# A novel preclinical mouse model recapitulates progressive phenotypes of Bryant-Li-Bhoj Syndrome

**DOI:** 10.64898/2026.06.16.732665

**Authors:** Dana E. Layo-Carris, Emily L. Durham, Emily E. Lubin, Annabel K. Sangree, Brianna Ciesielski, Marisol Hooks, Sarina M. Smith, Kaitlyn Worthington, Hannah M. Erdogan, Elizabeth M. Gonzalez, Xiao Min Wang, Erin E. Weiss, Kelly J. Abdalla, Divya Nair, W. Timothy O’Brien, Laura M. Bryant, Elizabeth J.K. Bhoj

## Abstract

Bryant-Li-Bhoj Syndrome (BLBS; OMIM: 619720, 619721) is a Mendelian neurogenetic condition, first described in 2020, with a mixed neurodevelopmental/neurodegenerative phenotype and variable systemic features. To date, 100 affected individuals with 74 unique causative variants have been published. Clinical data and prior functional work in multiple model systems have emphasized the utility of interrogating the pathogenesis of multiple causal variants to identify a convergent, therapeutically targetable mechanism. Additionally, the ability to evaluate the efficacy of future therapeutics relies on the availability of a robustly validated preclinical model. Here, we characterize the developmental and neurobehavioral phenotypes of a novel BLBS mouse model harboring one of the most recurrent causative variants (*h3-3a* p.T45I). H3.3^T45I^ mice recapitulate the BLBS natural history: perinatal growth restriction, delayed developmental milestones, and progressive motor and gait impairments. Adult mice additionally display craniofacial differences, impaired nest building, hyperactivity in a social context, and male-specific elevated aggression. The non-invasive, clinically translatable endpoints established here provide a validated preclinical platform for evaluating therapeutics for a community whose current standard of care is symptom management.

**Summary Statement:** A new mouse model mirrors the developmental delays, motor decline, and behavioral changes seen in individuals with this rare, progressive genetic brain disorder, providing a foundation for testing future therapies.

## Introduction

Pediatric neurodegenerative disorders are a devastating subset of Mendelian neurogenetic diagnoses that take an immeasurable toll on children, families, and society. Disease-modifying interventions are beginning to fundamentally alter the clinical trajectory for children living with these syndromes, including those diagnosed with spinal muscular atrophy, Duchenne muscular dystrophy, metachromatic leukodystrophy, cerebral adrenoleukodystrophy, and inborn errors of metabolism (Mendell et al., 2017; Mercuri et al., 2018; Mendell et al., 2025; Sabrina Haque et al., 2024; Gangji et al., 2025; Scott et al., 2023; Musunuru et al., 2025). Nonetheless, symptom management and palliative options are currently the standard of care for most affected individuals, especially those with ultra-rare or recently described neurogenetic conditions (Chan and Chang, 2024; Gunes et al., 2025).

A major barrier perpetuating the gap between the discovery of novel Mendelian disease genes and the generation of disease-modifying therapies, is the lack of validated model systems in which to perform mechanistic work and preclinical studies. Through this work, we seek to overcome that barrier for Bryant-Li-Bhoj Syndrome (BLBS; OMIM: 619720, 619721), a mixed neurodevelopmental and neurodegenerative disorder with variable multi-systemic effects that was first described in 2019 and is now reported to affect 100 people world-wide (Maver et al., 2019; Bryant et al., 2020; Okur et al., 2021; Khazaei et al., 2023; Layo-Carris et al., 2024; Hojo et al., 2024; Lubin et al., 2025; D’Onofrio et al., 2025). Heterozygous germline variants in either *H3-3A* (formerly *H3F3A*) or *H3-3B* (formerly *H3F3B*), the two genes that encode the H3.3 histone protein, cause BLBS. Regardless of causative variant, affected individuals most commonly exhibit developmental delay/intellectual disability, craniofacial anomalies, and tone differences that impact the neuromotor function over time (Okur et al., 2021; Layo-Carris et al., 2024). Notably, though, the clinical phenotype of affected individuals is highly variable, even amongst individuals sharing the same germline variant (Bryant et al., 2020; Layo-Carris et al., 2024). The phenotypic heterogeneity is not fully accounted for by sex, affected gene, or affected protein domain, and ongoing functional work seeks to more fully capture the molecular mechanisms underlying BLBS (Layo-Carris et al., 2024).

Before germline histone variants were identified as drivers of disease, histones were primarily known to play key roles in DNA packaging, gene regulation, structural support, and charge within cells (Vijayalakshmi et al., 2025; Maze et al., 2014). As reflected by the clinical phenotype of individuals with BLBS, who present with a mixed neurodevelopmental/neurodegenerative phenotype, as well in individuals with high grade gliomas, where somatic mutations in H3.3 are known drivers, H3.3 is crucial in development through its effect on stem cells and in postmitotic cells, such as neurons (Bryant et al., 2024). Knockdown of H3.3 in neural stem cells results in decreased proliferation and an early shift toward terminal differentiation in mouse brains (Xia and Jiao, 2017). Knockout of either *H3f3a* or *H3f3b,* the genes that encode for the murine H3.3 protein, has mixed results but the models generally show reduced viability and fertility when compared to unaffected littermates (Jang et al., 2015; Tang et al., 2015). Complete knockout of H3.3 results in embryonic lethality (Jang et al., 2015). Notably, the majority of research that has been done on the role of H3.3 in development relies on knockdown or knockout models, which is not a known mechanism of disease in humans. Though these models have been important for identifying the varied function of histones, the current models are insufficient to understand the human disease processes.

We and others have sought to address this translational gap. Single amino acid germline histone H3.3 substitutions modeled in mice (H3.3 p.G34R/V/W) have recapitulated the mixed neurodevelopmental and neurodegenerative features of BLBS, including phenotypic variability (Khazaei et al., 2023). This underscored the lesson originally demonstrated by affected individuals - there is significant phenotypic heterogeneity and, therefore, to best understand the shared therapeutically targetable point of mechanistic convergence, it is crucial to model a multitude of known disease-causing variants. This was further affirmed by subsequent data from the first 2D isogenic human induced pluripotent stem cell model and 3D dorsal forebrain organoids (H3.3 p.L48R) of BLBS (Sangree et al., 2026).

To continue building on this foundation of functional work towards validated preclinical models of and treatment for BLBS, we generated a novel BLBS mouse model that harbors the H3.3 p.T45I missense variant in *H3-3A* (Bramlage et al., 1997; Yuen and Knoepfler, 2013). Of the 72 unique causative BLBS variants, the p.T45I variant (often reported as p.T46I) is one of the most commonly occurring, shared by four unrelated individuals (Bryant et al., 2020; Layo-Carris et al., 2024). Here, we phenotypically and neurobehaviorally characterize the BLBS p.T45I mouse model informed by using clinically relevant assessments to evaluate its utility as a preclinical model system. Our robust interrogation allows us to establish non-invasive metrics that can be easily quantified in the evaluation of novel therapeutics for a community whose current standard of care is symptom management.

## Results

### Gross assessment of perinatal murine phenotypes

The heterozygous germline *H3-3A* p.T45I missense variant is one of the most commonly represented variants across the BLBS community (Figure 1A). It is shared by four unrelated individuals, whose phenotypes are described in Table 1. This variant was used to create a novel murine model of BLBS.

**Figure 1.**
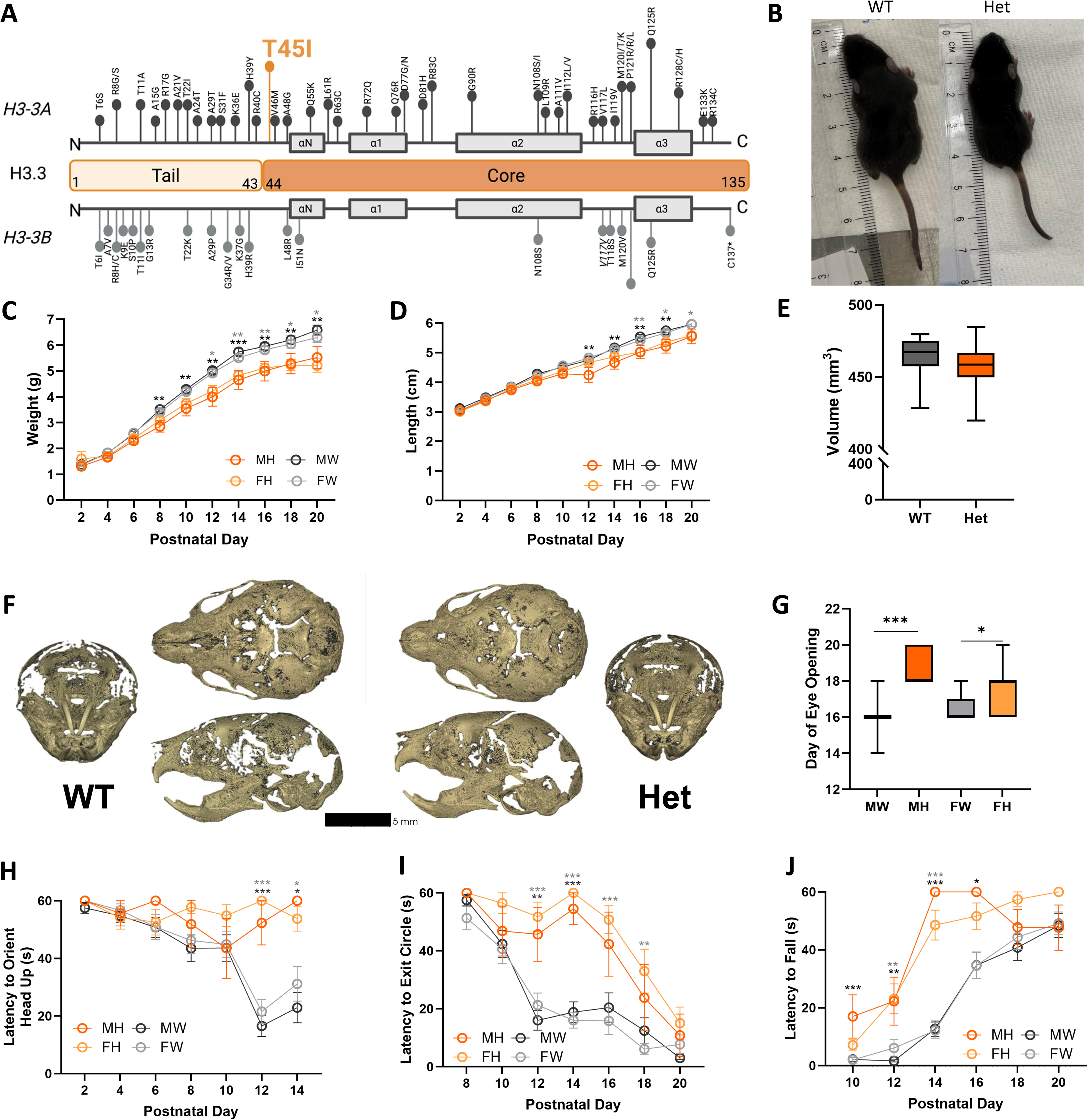
BLBS p.T45I mice show delayed developmental and neuromuscular reflex milestones. A. Graphic of all reported BLBS variants reported to date mapped to gene of origin (*H3-3A* in charcoal on top or *H3-3B* in light gray on bottom) and to the H3.3 protein (divided into tail and core domains). Length of lollipop corresponds to number of affected individuals. This mouse model harbors the *H3-*3A c.137C>T, P.T45I variant. Adapted from (Layo-Carris et al., 2024). B. Image of H3.3^WT^ (left) and H3.3^T45I^ (right) mouse for gross phenotypic comparison. C-J. Generally, H3.3^WT^ data is presented in shades of grey and H3.3^T45I^ data is presented in shades of orange. Specifically, male H3.3^WT^ data is black/dark gray, female H3.3^WT^ data is light grey, male H3.3^T45I^ data is orange, and female H3.3^T45I^ data is pale orange. C. H3.3^T45I^ pups weigh significantly less than H3.3^WT^ littermates starting at PND 8. D. H3.3^T45I^ pups show a statistically significant reduction in length compared to H3.3^WT^ littermates starting at PND 12. E. Endocast volume is not significantly different between genotypes, and skull shape is grossly similar. F. Representative 3D Micro-CT renderings of P0 H3.3^WT^ (left) and H3.3^T45I^ (right) displayed with anterior views (lateral) and superior (top) and left lateral (bottom) views centrally. n = 4 per genotype. G. Both male and female H3.3^T45I^ pups had delayed eye opening compared to H3.3^WT^ littermates. H. Both male and female H3.3^T45I^ pups showed statistically significant delayed latency to orient head up in the negative geotaxis assay compared to H3.3^WT^ littermates beginning at PND 12. I. Both male and female H3.3^T45I^ pups showed statistically significant delayed latency to exit circle compared to H3.3^WT^ littermates beginning at PND 12 and continuing through PND 18. J. Both male and female H3.3^T45I^ pups showed a significantly larger latency to fall compared to H3.3^WT^ littermates from PND 10 - PND 16. n ≥ 8 for assessments other than Micro-CT. MW = male WT; MH = male Het; FW = female WT; FH = female Het. *p≤0.05, ** p≤0.01 *** p≤0.001. Scale indicated on figure.

**Table 1.**
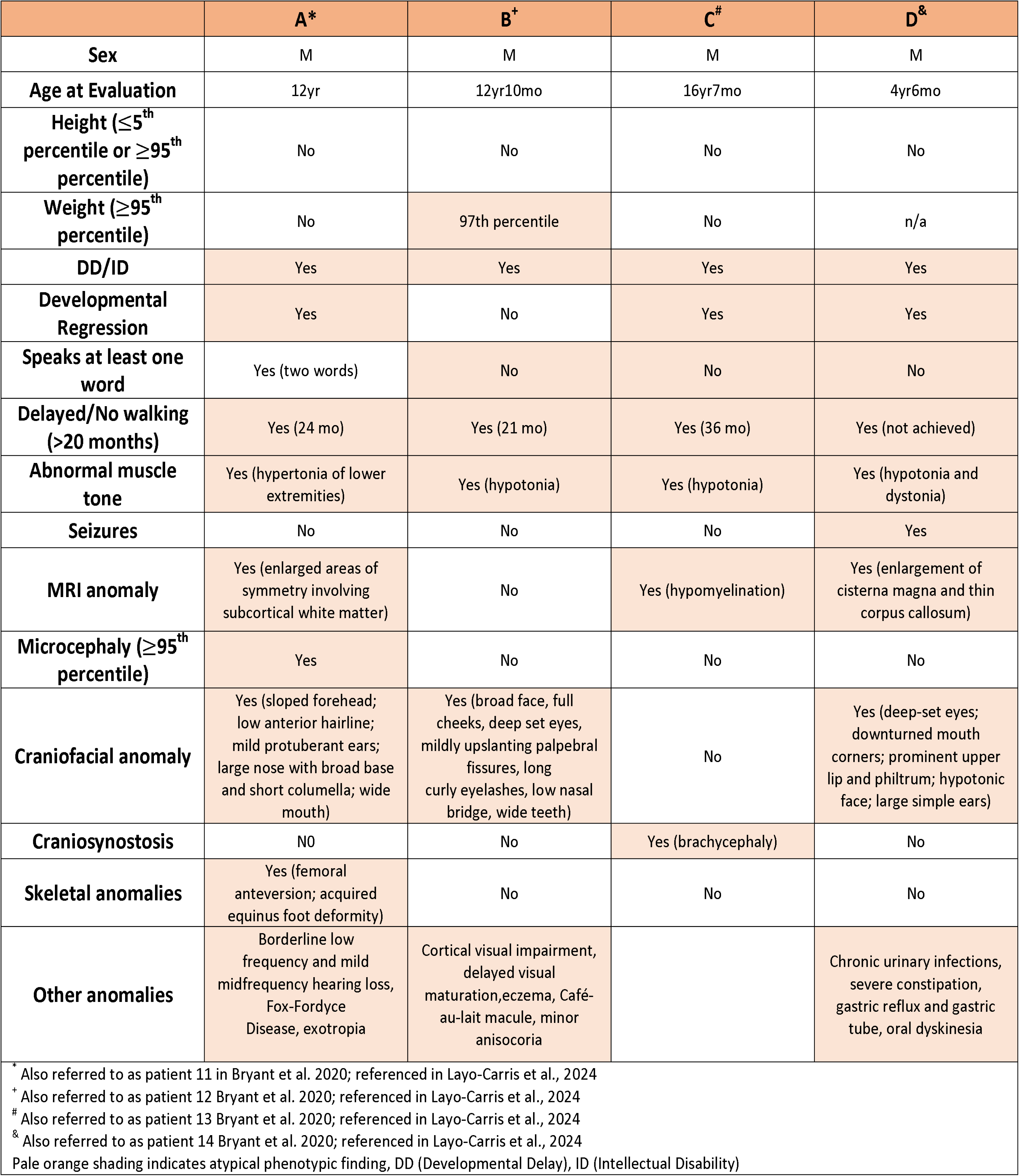
BLBS H3F3A c.137C>T p.T45I phenotype.

|  | A* | B<sup>+</sup> | C<sup>#</sup> | D<sup>&</sup> |
| --- | --- | --- | --- | --- |
| Sex | M | M | M | M |
| Age at Evaluation | 12yr | 12yr10mo | 16yr7mo | 4yr6mo |
| Height ($\leq 5^{\text{th}}$ percentile or $\geq 95^{\text{th}}$ percentile) | No | No | No | No |
| Weight ($\geq 95^{\text{th}}$ percentile) | No | 97th percentile | No | n/a |
| DD/ID | Yes | Yes | Yes | Yes |
| Developmental Regression | Yes | No | Yes | Yes |
| Speaks at least one word | Yes (two words) | No | No | No |
| Delayed/No walking (>20 months) | Yes (24 mo) | Yes (21 mo) | Yes (36 mo) | Yes (not achieved) |
| Abnormal muscle tone | Yes (hypertonia of lower extremities) | Yes (hypotonia) | Yes (hypotonia) | Yes (hypotonia and dystonia) |
| Seizures | No | No | No | Yes |
| MRI anomaly | Yes (enlarged areas of symmetry involving subcortical white matter) | No | Yes (hypomyelination) | Yes (enlargement of cisterna magna and thin corpus callosum) |
| Microcephaly ($\geq 95^{\text{th}}$ percentile) | Yes | No | No | No |
| Craniofacial anomaly | Yes (sloped forehead; low anterior hairline; mild protuberant ears; large nose with broad base and short columella; wide mouth) | Yes (broad face, full cheeks, deep set eyes, mildly upslanting palpebral fissures, long curly eyelashes, low nasal bridge, wide teeth) | No | Yes (deep-set eyes; downturned mouth corners; prominent upper lip and philtrum; hypotonic face; large simple ears) |
| Craniosynostosis | NO | No | Yes (brachycephaly) | No |
| Skeletal anomalies | Yes (femoral anteversion; acquired equinus foot deformity) | No | No | No |
| Other anomalies | Borderline low frequency and mild midfrequency hearing loss, Fox-Fordyce Disease, exotropia | Cortical visual impairment, delayed visual maturation,eczema, Café-au-lait macule, minor anisocoria |  | Chronic urinary infections, severe constipation, gastric reflux and gastric tube, oral dyskinesia |
| * Also referred to as patient 11 in Bryant et al. 2020; referenced in Layo-Carris et al., 2024<br><sup>+</sup> Also referred to as patient 12 Bryant et al. 2020; referenced in Layo-Carris et al., 2024<br><sup>#</sup> Also referred to as patient 13 Bryant et al. 2020; referenced in Layo-Carris et al., 2024<br><sup>&</sup> Also referred to as patient 14 Bryant et al. 2020; referenced in Layo-Carris et al., 2024<br>Pale orange shading indicates atypical phenotypic finding, DD (Developmental Delay), ID (Intellectual Disability) |  |  |  |  |

We first performed a comprehensive assessment of gross perinatal mouse phenotypes. Prior studies repeatedly demonstrate that homozygous null H3.3 variants are incompatible with life, and all patients are heterozygous, developmental milestones were assessed in at least 10 litters of pups, which were comprised of heterozygous H3.3^T45I^ mice and H3.3^WT^ littermates. Assessments occurred every other day between post-natal day 2 (PND 2) and PND 20. Gross phenotypic differences were apparent (Figure 1B). While weight is equivalent between genotypes at PND 2, a genotype- and sex-based dichotomy emerged quicky (Figure 1C). Male H3.3^T45I^ pup weight diverged as significantly less than their littermates at PND 8 (p=0.002) and the female H3.3^T45I^ pup weight diverged as significantly less at PND 12 (p=0.011). The statistically significant weight differences persisted through PND 20 (PND 10: males = 0.007; PND 12: males = 0.002; PND 14: males ≤ 0.001, females = 0.005; PND 16: males = 0.002, females = 0.006; PND 18: males = 0.013, females = 0.007; PND20: males p=0.02, females p=0.003). A similar trend was observed for length (Figure 1D). The lengths of the H3.3^T45I^ pups are significantly smaller than their unaffected littermates beginning at PND 12 (PND 12: males p=0.003) and continuing through PND 20 (PND 14: males p=0.004; PND 16: males p=0.008, females p=0.008; PND 18: males p=0.004, females p=0.027; PND 20: females p=0.016). Evaluation of space occupied by the brain at P0 shows no difference between H3.3^WT^ and H3.3^T45I^ pups (p=0.222) (Figure 1E-F). No differences in the gross anatomy of skulls were identified between genotypes indicating that differences potentially occur later in development (Supplemental Figure 1A). H3.3^T45I^ pups had delayed eye opening when compared to unaffected littermates (male H3.3^WT^ vs male H3.3^T45I^: p≤0.001; male H3.3^WT^ vs female H3.3^T45I^: p=0.006; female H3.3^WT^ vs male H3.3^T45I^: p≤0.001; female H3.3^WT^ vs female H3.3^T45I^: p=0.046) (Figure 1G). No differences were seen between H3.3^T45I^ pups and unaffected littermates in fur growth, pinnae detachment, or incisor eruption (Supplemental Figure 1B-D).

Perinatal pre-weaned mice were also assessed with a number of neuromuscular reflexes and simple behavioral tests. In assays that interrogated motor coordination, male and female H3.3^T45I^ pups demonstrated both significantly delayed negative geotaxis (Figure 1H) and significantly delayed latency to exit circle (Figure 1I) beginning at PND 12 when compared to unaffected littermates (Geotaxis – PND12: males p<0.001, females p<0.001; PND14: males p=0.049, females p=0.03; Circle – PND12: males p=0.002, females p≤0.001; PND14: males p≤0.001, females p≤0.001; PND16: females p≤0.001; PND18: females p=0.003). Additionally, as a measure of grip strength (vertical wire hang), H3.3^T45I^ pups showed a significantly increased latency to fall beginning at PND 10 as compared to unaffected littermates (PND10: males p≤0.001; PND12: males p=0.003, females p=0.003; PND14: males p≤0.001, females p≤0.001; PND16: males p=0.029) (Figure 1J),however this may be due to the significantly smaller weight of H3.3^T45I^ pups (Figure 1C). No differences were seen between H3.3^T45I^ pups and unaffected littermates in the appearance of the grasping reflex or latency to prone position in surface righting assay (Supplemental Figure 1E-F).

### Longitudinal assessment of motor development

Since individuals with BLBS exhibit a progressive neuromotor phenotype as they age, including the development of ataxic gait, the motor function of H3.3^T45I^ was longitudinally assessed compared to unaffected littermates in both adult mice 12-16 weeks (Figure 2) and in aged mice >60 weeks (Figure 3).

**Figure 2.**
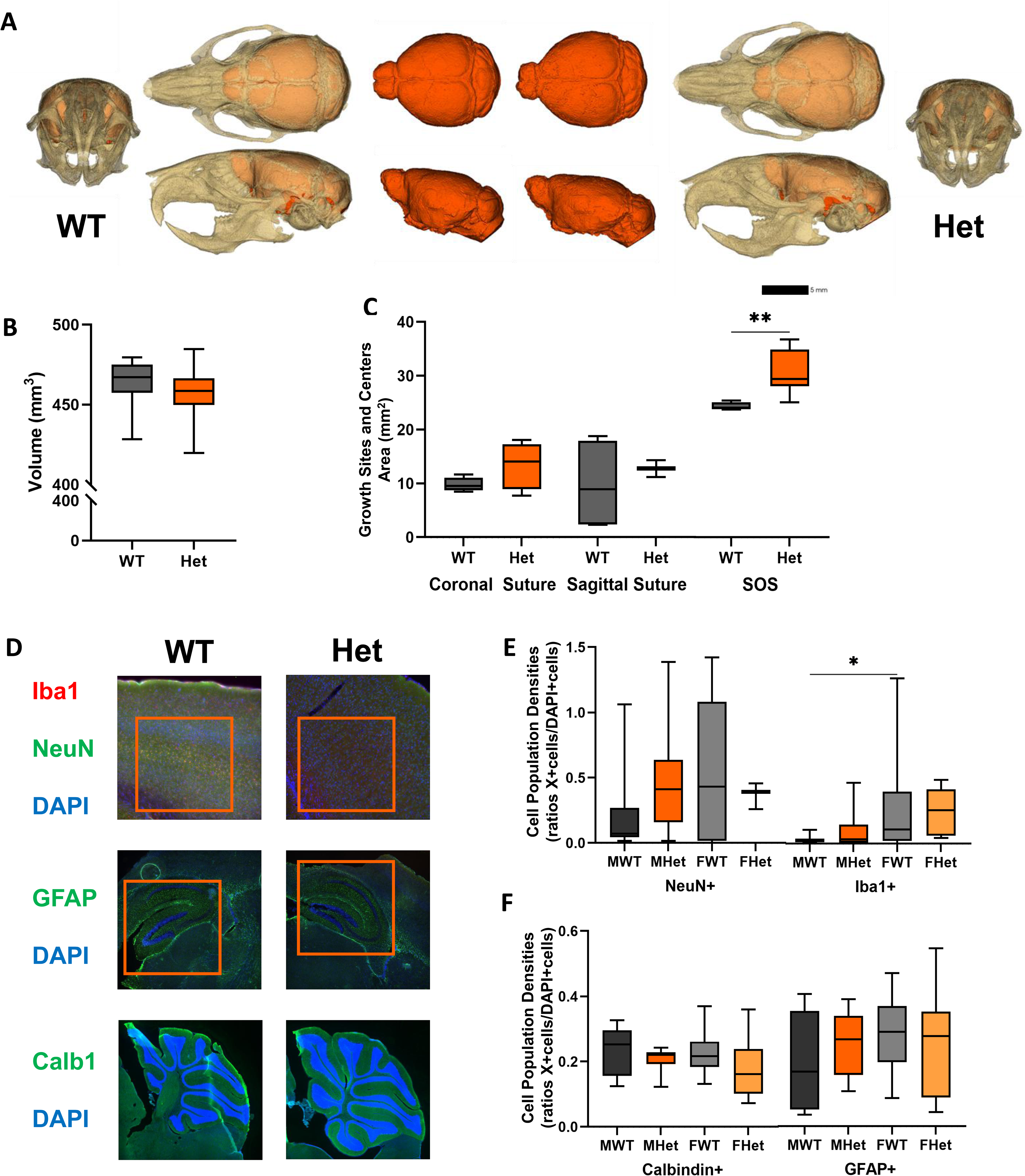
Adult BLBS p.T45I mice show altered skull growth and brain cell composition. Generally, H3.3^WT^ data is presented in shades of grey and H3.3^T45I^ data is presented in shades of orange. Specifically, male H3.3^WT^ data is black/dark gray, female H3.3^WT^ data is light grey, male H3.3^T45I^ data is orange, and female H3.3^T45I^ data is pale orange. A. Representative 3D micro-CT renderings of adult H3.3^WT^ (left) and H3.3^T45I^ (right) displayed with anterior views (lateral) and superior (top) and left lateral (bottom) views with isolated endocasts displayed centrally. B-C. Endocast volume assessment indicates similar braincase size between genotypes however, growth centers (SOS) are altered in H3.3^T45I^ as compared to H3.3^WT^ littermates. D-F. Representative immunohistochemical images at 20x magnification with standard ROI identified (orange box) for H3.3^WT^ (left) and H3.3^T45I^ (right) mouse brain sections. Targets are indicated by color in the left margin and quantification of target cells standardized to number of cells within the region of interest is displayed in E-F. n ≥ 8 MW = male WT; MH = male Het; FW = female WT; FH = female Het; SOS = Spheno-occipital Synchondrosis. *p≤0.05, ** p≤0.01. Scale indicated on figure.

**Figure 3.**
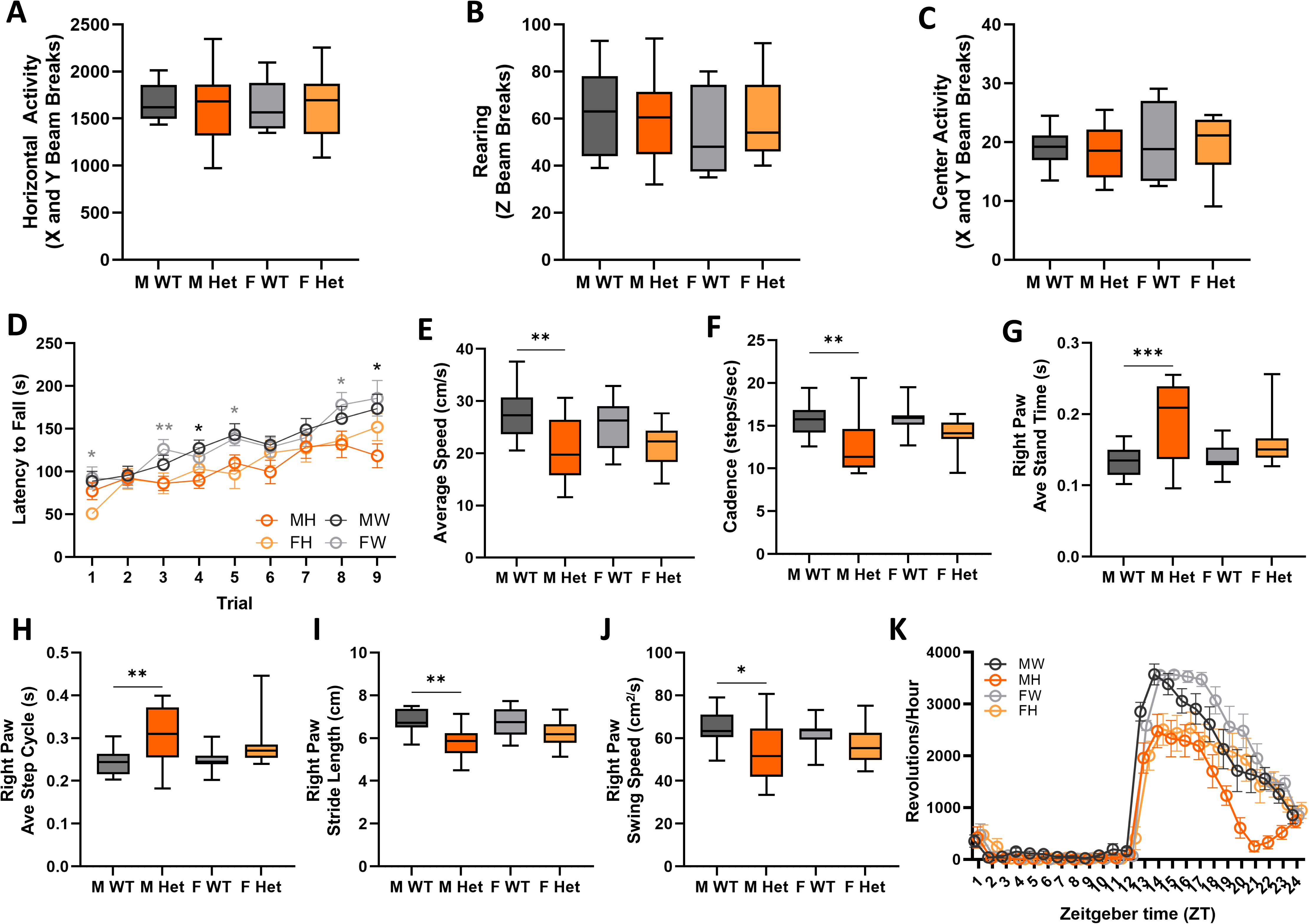
Adult BLBS p.T45I mice show deficits in gross motor and gait that are more profound in males. A-K. Male H3.3^WT^ data is black/dark gray, female H3.3^WT^ data is light grey, male H3.3^T45I^ data is orange, and female H3.3^T45I^ data is pale orange. A-C. Open field assay. A. No observed differences between H3.3^T45I^ mice and H3.3^WT^ littermates in horizontal/general ambulatory activity. B. No observed differences between H3.3^T45I^ mice and H3.3^WT^ littermates in rearing activity. C. No observed differences between H3.3^T45I^ mice and H3.3^WT^ littermates in center activity. D. Over three days of accelerating rotarod (3 trials per day), H3.3^T45I^ mice are variably worse than H3.3^WT^ littermates in gross motor coordination, with decreased latency to fall. E-J. Noldus CatWalkXT System data. E. Male H3.3^T45I^ mice show statistically significant decrease in average speed than H3.3^WT^ littermates. F. Male H3.3^T45I^ mice show statistically significant decrease in cadence when compared to H3.3^WT^ littermates. G. Male H3.3^T45I^ mice show statistically significant increase in right paw average stand time when compared to H3.3^WT^ littermates. H. Male H3.3^T45I^ mice show statistically significant increase in right paw average stand cycle when compared to H3.3^WT^ littermates. I. Male H3.3^T45I^ mice show statistically significant decrease in right paw stride length when compared to H3.3^WT^ littermates. J. Male H3.3^T45I^ mice show statistically significant decrease in right paw swing speed when compared to H3.3^WT^ littermates. K. No significant difference is seen in endogenous period (tau), H3.3^T45I^ mice show a significant reduction of running wheel activity. Reduced activity is seen more so in males during ZT13-17 and females later in the dark phase ZT21-23. n ≥ 8 MW = male WT; MH = male Het; FW = female WT; FH = female Het. *p≤0.05, ** p≤0.01 *** p≤0.001.

### Gross assessment of adult murine phenotypes

Gross differences in pups are subtle between PND 10-20 (Figure 1B) which continues throughout life. By adulthood, assessment of the space occupied by the brain and metrics of craniofacial form indicated no statistically significant morphological changes between genotypes (Figure 2A, 2B). Still, skulls of H3.3^T45I^ mice appear to be longer and less domed as compared to their unaffected littermates (Supplemental Figure 2M). This indicates that both brain and skull may be affected by alterations in H3.3. Investigation of growth sites (sutures) and centers (SOS) revealed that H3.3^T45I^ mice have a larger growth center as compared to H3.3^WT^ littermates (Figure 2C, Supplemental Figure 2N).

Similarly, investigation of cell types in the brain using immunohistochemistry indicated few differences by genotype in the number of neurons (NeuN, p=0.559), astrocytes (GFAP, p=0.4742), and Purkinje cells (Calbindin, p=0.105) when signal counts were standardized by total cell count within the indicated region of interest (Figure 2D-F). Iba1+ microglia were similar between genotypes (p=0.756) but there were significantly more microglia in female H3.3^WT^ brains as compared to their male H3.3^WT^ littermates (p=0.029) (Figure 2E).

### Adult Mice - Motor

In addition to craniofacial morphometrics, adult mice were assessed for spontaneous activity in an open field (Figures 3A-3C). This assay allows for the interrogation of ambulation, rearing activity (Figure 3A, 3B) and anxiety-related phenotypes (Figure 3C). No differences were seen between H3.3^T45I^ mice and their unaffected littermates in ambulatory activity (Figure 3A), rearing (Figure 3B), or in center activity (Figure 3C). To evaluate balance/coordination and motor learning, the accelerating rotarod was employed (Figure 3D). Throughout 9 trials (3 trials per day over 3 consecutive days), H3.3^T45I^ mice present a mild, but statistically significant, deficit in balance/coordination. The subtle deficit is underscored by only a few trials where the H3.3^T45I^ mice have a decreased latency to fall off the rotarod when compared to unaffected littermates (Trial 1: females p=0.011; Trial 3: females p=0.006; Trial 4: males p=0.026; Trial 5: females p=0.015; Trial 8: females p=0.027; Trial 9: males p=0.021).

To further quantitatively dissect motor output, gait was assessed by the Noldus CatWalkXT, which measures over 180 parameters of gait and paw dynamics (Supplemental Figure 2). Gait analysis showed decreased speed of ambulation, as well as cadence in the H3.3^T45I^ male mice compared to unaffected male littermates (Speed: p=0.01; Cadence: p=0.006) (Figures 3E-F). Analysis of paw dynamics showed that male H3.3^T45I^ mice have increased stand time (Stand Time: p=0.006). All gait and paw metrics are included in Supplemental Figure 2. Further, a histological assessment indicated subtle differences in muscle fibers in the lower limbs (Supplemental Figure 2O-P). Interestingly, unlike H3.3^T45I^ male mice, H3.3^T45I^ female mice do not show statistically significant fine motor phenotypes compared to unaffected littermates. Based on clinical data, this lack of profoundly impaired motor function in all H3.3^T45I^ mice, even with this granular analysis, was unexpected however this may be related to the quadrupedal nature of the mouse.

Finally, wheel running activity was used to investigate circadian rhythm and voluntary motor activity that may be related to early motor impairment. Data was acquired over several days in a standard light cycle (ZT1-ZT12 is the “lights on” phaseandZT13-ZT24 are the “lights off” phase). During the last 14 days of the procedure, the lighting schedule was switched to complete darkness to assess the endogenous period of activity. There were no differences between the genotypes in phase shift or endogenous period (tau). However, a significant reduction in wheel running activity and pattern of activity was seen in the H3.3^T45I^ mice when compared to H3.3^WT^ mice (ZT13: males p=0.036; ZT14: males p=0.013, females p=0.008; ZT15: males p=0.034, females p=0.009; ZT16: females p=0.024; ZT17 females p=0.011; ZT21: males p=0.028; ZT22: males p=0.002; ZT23: males p=0.024) (Figure 3K). At the start of the mouse wake cycle (ZT12), there is increased wheel rotations per hour for all mice. However, the H3.3^T45I^ mice plateau at rates significantly below unaffected littermates between ZT13-16, with the H3.3^T45I^ females steadily maintaining activity levels and the males showing a statistically significant decrease in activity between ZT19-23. These findings may be confounded by the irregular gait in the H3.3^T45I^ males demonstrated in the CatWalk data (Figures 3E-3J), which may make running on a slotted wheel difficult. General activity on the floor of the wheel running chamber was not recorded. This sex-based dichotomy in the mouse model was notable since this trend was not observed in BLBS affected individuals (Okur et al., 2021).

### Aged mice

Because the limited longitudinal data available for individuals living with BLBS shows that progressive motor decline worsens with age, we re-evaluated the gross and fine motor coordination of elder H3.3^T45I^ mice using an abridged behavioral battery with the rotarod and CatWalkXT system.

Notably, 60 weeks of age in mice roughly correlates with 40-60 years of age in humans, which is about the age of the oldest published evaluation of an individual living with BLBS. The rotarod data show motor impairment in the H3.3^T45I^ mice, with significantly decreased latency to fall over all nine trials (Trial 1: males p≤0.001, females p≤0.001; Trial 2: males p=0.001; Trial 3: males p=0.003, females p≤0.001; Trial 4: males p=0.001, females p=0.01; Trial 5: males p≤0.001, females p≤0.001; Trial 6: males p≤0.001, females p=0.001; Trial 7: males p=0.002, females p≤0.001; Trial 8: females p=0.003; Trial 9: males p=0.018, females p≤0.001) (Figure 4A). The gait analysis showed a larger variation within H3.3^T45I^ mice groups, with overall less significant measurements than at the adult time point (Stride Length: males p≤0.001, females p=0.028) (Figure 4B-G). While only right paw data are shown in Figure 3, all metrics are included in Supplemental Figure 4. Interestingly, unlike in the adult mouse cohort, no sex-driven differences were observed.

**Figure 4.**
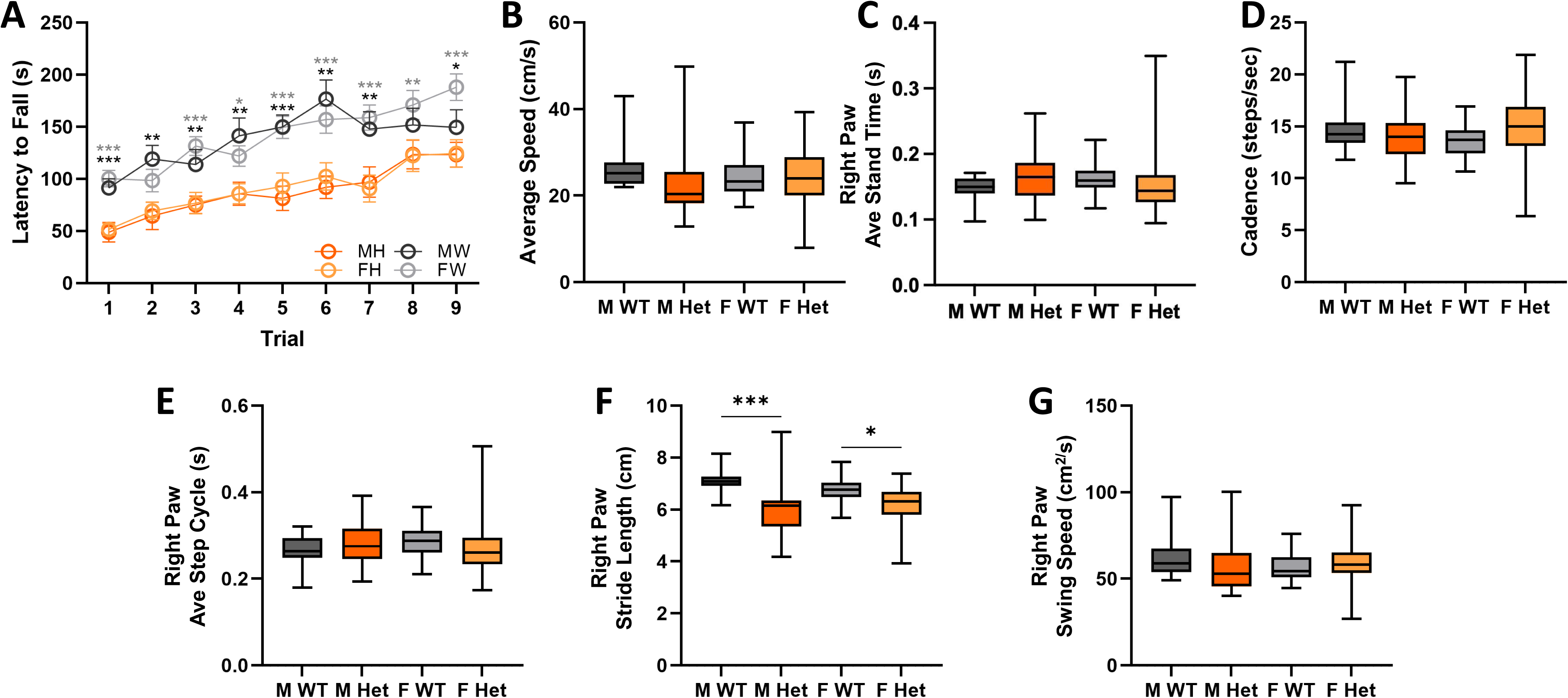
Aged BLBS p.T45I mice show sex-agnostic motor deficits. A-G. Male H3.3^WT^ data is black/dark gray, female H3.3^WT^ data is light grey, male H3.3^T45I^ data is orange, and female H3.3^T45I^ data is pale orange. A. H3.3^T45I^ mice show significantly decreased latency to fall over 9 trials (3 trials per day over 3 consecutive days) on rotarod compared to H3.3^WT^ littermates. B-G. Noldus CatWalkXT data. B. No significant differences in average speed between H3.3^T45I^ mice and H3.3^WT^ littermates. C. No significant differences in cadence between H3.3^T45I^ mice and H3.3^WT^ littermates. D. No significant differences in right paw stand time between H3.3^T45I^ mice and H3.3^WT^ littermates. E. No significant differences in paw step cycle between H3.3^T45I^ mice and H3.3^WT^ littermates. F. H3.3^T45I^ mice show statistically significant decrease in stride length compared to H3.3^WT^ littermates. G. No significant differences in paw swing speed between H3.3^T45I^ mice and H3.3^WT^ littermates. n ≥ 8 MW = male WT; MH = male Het; FW = female WT; FH = female Het. *p≤0.05, ** p≤0.01 *** p≤0.001.

### Comprehensive assessment of communication, social, and cognitive development

Clinically relevant developmental domains beyond gross and fine motor function include communication, social, and cognitive development, which are known to be impacted in individuals with BLBS, including those harboring the p.T45I variant (Table 1) (Layo-Carris et al., 2024). For example, 3 of the 4 reported individuals with this variant have phenotypic features that could impact expressive or receptive communication. Thus, we assessed communication with ultra-sonic vocalizations (USV) in pre-weaned pups (Supplemental Figure 5). While male and female H3.3^T45I^ mice showed USV call numbers and durations that were less than their unaffected littermates, there was not a statistically significant difference (Supplemental Figures 5A- B). There were no clear sex- or genotype differences in USV frequency (Supplemental Figure 5C).

Further, individuals with BLBS are often diagnosed with developmental delay/intellectual disability, anxiety, and/or autism spectrum disorder. To evaluate if this novel BLBS model recapitulated these clinically relevant phenotypes, we conducted assays that measure social behavior and cognition of the adult H3.3^T45I^ mouse line. In elevated zero maze, another test to measure anxiety-related behavior, data show no difference in latency to exit between H3.3^T45I^ mice and controls (Figure 5A) and no difference in preference to remain in the closed arm of the maze when comparing H3.3^T45I^ and H3.3^WT^ mice (Figure 5B). A nest building assay was used to measure obsessive compulsive-like behaviors and showed that, while H3.3^WT^ mice demonstrated significant improvement in nest scores over the 24-hour period, H3.3^T45I^ mice showed no improvement over the same time period, often not building a complete nest (between time point: male H3.3^WT^ p=0.012, female H3.3^WT^ p=0.021; score change at 24 hours: males p=0.006, females p=0.002) (Figure 5C).

**Figure 5.**
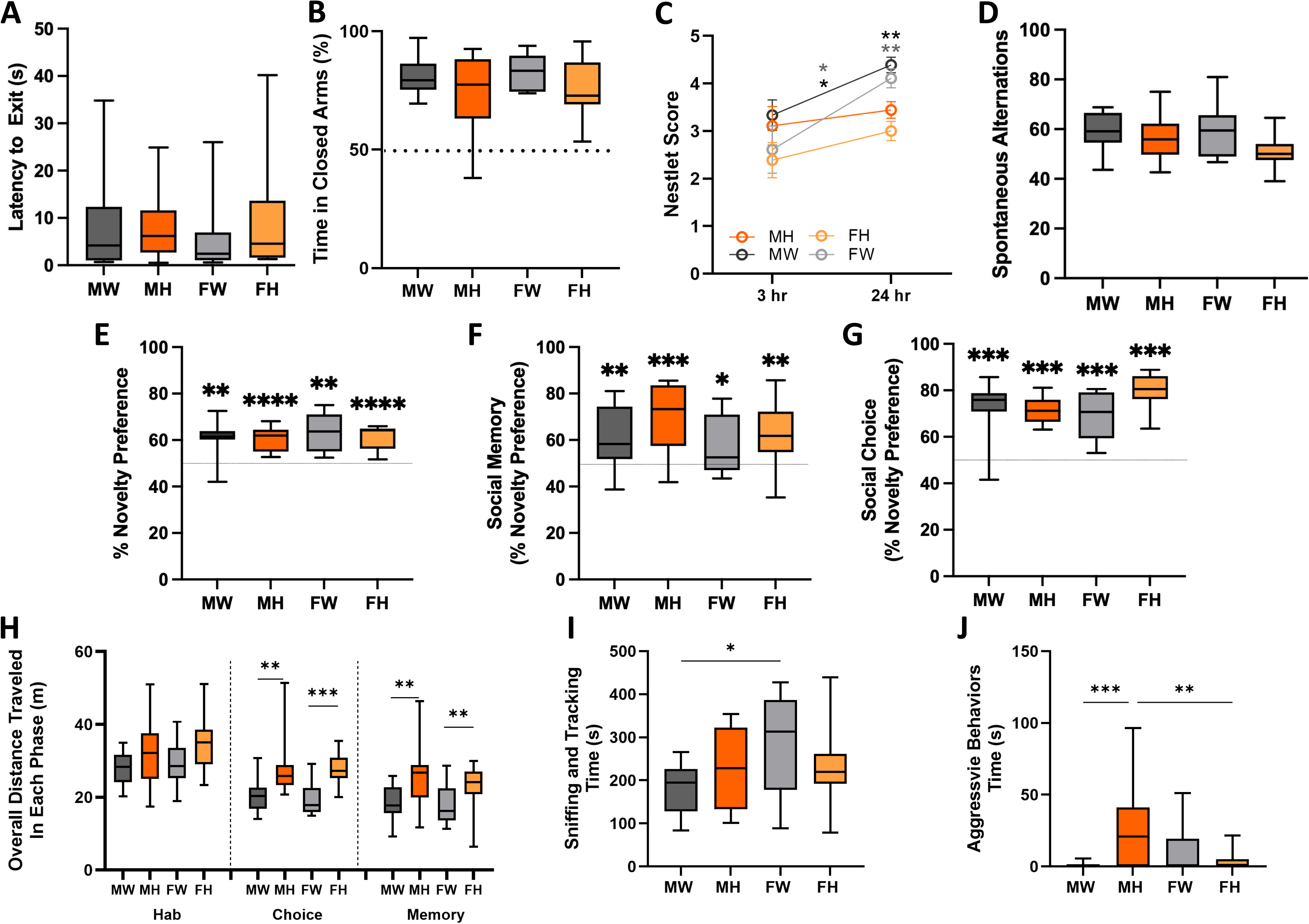
Social and cognitive development impacted in adult BLBS p.T45I mice. A-J. Male H3.3^WT^ data is black/dark gray, female H3.3^WT^ data is light grey, male H3.3^T45I^ data is orange, and female H3.3^T45I^ data is pale orange. A. No significant differences in latency to exit between H3.3^T45I^ mice and H3.3^WT^ littermates in elevated zero maze. B. No significant differences in time in closed arms of elevated zero maze between H3.3^T45I^ mice and H3.3^WT^ littermates. C. In the nest building assay, H3.3^WT^ mice demonstrated significant improvement in nest scores over the 24-hour period while H3.3^T45I^ mice showed no improvement over the same time period. D. No significant differences in time in spontaneous alternations between H3.3^T45I^ mice and H3.3^WT^ littermates. E. All mice, regardless of sex or genotype, showed novelty preference in novel object recognition assay. F-H. Three-chamber social choice and memory task. F. All mice, regardless of sex or genotype, showed preference for mouse over rock stimuli. G. All mice, regardless of sex or genotype, showed preference for novel mouse over familiar mouse. H. Male and female H3.3^T45I^ mice traveled significantly more distance in the choice and memory phases compared to H3.3^WT^ littermates. I. Female H3.3^WT^ mice spent significantly more time sniffing and tracking as compared to their male H3.3^WT^ counterparts. J. Male H3.3^T45I^ mice were significantly more aggressive than male H3.3^WT^ mice in the resident intruder assay. n ≥ 8 MW = male WT; MH = male Het; FW = female WT; FH = female Het. *p≤0.05, ** p≤0.01 *** p≤0.001.

To assess working memory, spontaneous alternations in a Y-maze was assessed. There was no difference between groups for percent of spontaneous alternations suggesting that working memory was intact. Long-term memory was assessed with the Novel Object Recognition (NOR) procedure. In NOR, the mice are familiarized with a pair of similar objects. Twenty-four hours later one of the paired objects is replaced with a novel object. If familiarization occurred during training, mice preferentially explore the novel object. A failure to do so during recall suggests failure to recall familiarity of the paired objects during training. All groups demonstrated novelty preference in the NOR test (male H3.3^WT^ p=0.01, male H3.3^T45I^ p≤0.001, female H3.3^WT^ p=0.002, female H3.3^T45I^ p≤0.001) (Figure 5D and E). To test social behavior, the three-chamber social choice procedure was performed with an additional social memory phase. When mice are provided the opportunity to explore an object (inanimate cue) and a novel mouse (social cue), mice innately show a preference for exploration of the social cue . In the social preference phase, all mice showed preference for the social cue over the inanimate cue (male H3.3^WT^ p≤0.001, male H3.3^T45I^ p≤0.001, female H3.3^WT^ p≤0.001, female H3.3^T45I^ p≤0.001) (Figure 5F).

Immediately after the social choice phase of the three-chamber procedure, the experimental mouse is tested with a new mouse that replaced the inanimate object. Mice preferentially explore the novel mouse over the familiarized mouse. All groups showed preference for the novel mouse (male H3.3^WT^ p=0.005, male H3.3^T45I^ p≤0.001, female H3.3^WT^ p=0.029, female H3.3^T45I^ p=0.002) (Figure 5G). While there were not sex- or genotype-based changes observed in the quantified metrics, the H3.3^T45I^ mice were noted to be comparatively hyperactive in the assay. We found that they traveled significantly more distance than unaffected littermates in the social phases of the task (Choice: males p=0.002, females p≤0.001; Memory: males p=0.01, females p=0.008) (Figure 5H). It was also noted that male H3.3^T45I^ mice seemed to be investigating the social stimuli differently than the H3.3^WT^ mice. When quantified, the male H3.3^T45I^ mice were found to spend more time rearing against or mounting the container housing the social stimuli during investigation while H3.3^WT^ mice merely sniffed and nose-poked for investigation (Figure 5I). This heightened exploration suggested that there may be an antisocial or aggressive phenotype in the mice. We, therefore, performed a resident-intruder assay, in which H3.3^T45I^ mice were singly housed for 10 days before a novel “intruder” mouse was introduced. Behavior was then monitored for sniffing/tracking events as well as mounting/fighting behaviors for 10 minutes. If the fighting became serious, the assay was stopped and the intruder mouse was removed. Of the 14 H3.3^T45I^ male mice tested, 5 of the sessions had to be ended early due to overly aggressive interactions. Overall, the male H3.3^T45I^ mice were significantly more aggressive than male H3.3^WT^ mice, whereas females showed no aggressive phenotype (male H3.3^WT^ vs male H3.3^T45I^ p≤0.001, male H3.3^T45I^ vs female H3.3^T45I^ p≤0.001) (Figure 5J).

## Discussion

More than 300 million people worldwide live with a rare disease, and 95% of them lack targeted, disease-modifying therapeutic interventions (The Lancet Global Health, 2024). Staggeringly, in high-income countries, a comparable number of children with pediatric neurodegenerative conditions and pediatric cancers die each year but, due in part to disparities in awareness and resource allocation, there is a notable dichotomy in available treatments for these two populations (Elvidge et al., 2025). This results in both tremendous psychosocial impact on families navigating rare disease diagnostic odysseys with uncertain clinical management pathways and on society due to health care utilization (Navarrete-Opazo et al., 2021; Nevin et al., 2023). Nonetheless, the field of pediatric oncology demonstrates that progress is possible, including for children with H3.3 somatic-variant-driven malignancies (Elvidge et al., 2025; Helms et al., 2023; Monje et al., 2025). Contributing to this chasm between diagnosis and therapeutic development for individuals living with rare genetic conditions is the fact that the pace of Mendelian disease gene discovery surpasses our ability to develop preclinical model systems in which to perform functional validation. Here, we address the current lack of a putative preclinical mouse model of BLBS that is characterized in a way that is informed by the known progressive phenotype of this condition.

Prior reports have demonstrated that BLBS is a mixed neurodevelopmental/neurodegenerative condition in which affected individuals commonly exhibit gross phenotypic differences in growth (weight and length), craniofacial form, attainment of developmental milestones, and neuromotor function that seems to worsen as the affected individual ages (Maver et al., 2019; Bryant et al., 2020; Okur et al., 2021; Khazaei et al., 2023; Layo-Carris et al., 2024; Hojo et al., 2024; Lubin et al., 2025; D’Onofrio et al., 2025). Fortunately, growth, craniofacial development, and attainment of developmental milestones are features that are non-invasively assessed beginning at a child’s first visit to their pediatrician and reassessed at every subsequent visit. These traits, in conjunction with neuromuscular function, have a significant impact on quality of life. The FDA has recently developed a series on patient-focused drug development guidance that emphasize the patient and caregiver experience at the center of drug development and regulatory decision making (FDA Center for Drug Evaluation and Research, 2018; FDA Center for Drug Evaluation and Research, 2022; FDA Center for Drug Evaluation and Research et al., 2025; FDA Center for Drug Evaluation and Research et al., 2023). Thus, understanding what aspects of disease pathogenesis are most prominent for affected individuals, and then being able to quantify the efficacy of a novel intervention in addressing these, is fundamental for translational research efforts. It is also essential that these patient-relevant phenotypes are recapitulated in any preclinical model system being used to evaluate a putative disease-modifying intervention.

Thus, as part of our comprehensive phenotyping of this novel BLBS p.T45I mouse model, we assessed patient-relevant features (Motch Perrine et al., 2021; Motch Perrine et al., 2023; Mitteroecker and Stansfield, 2021). Mirroring the BLBS natural history observed in affected individuals harboring this common variant, we found that from the earliest developmental stages and continuing through their lifetime, H3.3^T45I^ mice showed statistically significant differences in growth (Figure 1), craniofacial development (Figure 1, Figure 2, Supplemental Figures 1-2), attainment of developmental milestones (Figure 1, Figure 3), and progressive decline in neuromuscular function compared to H3.3^WT^ littermates (Figures 3-5). Reduced body weight and length emerged as early as PND 8 and PND 12, respectively, consistent with the growth delays reported across the broader BLBS cohort (Layo-Carris et al., 2024).

Delayed eye opening, a murine correlate of sensory maturation, further reflected the developmental-delay characteristic of the syndrome. Importantly, assessment of the space occupied by the brain at P0 revealed no statistically significant difference in endocranial volume between H3.3^T45I^ and H3.3^WT^ pups (Figure 1E, 1F), indicating that craniofacial and brain morphology differences in BLBS potentially arise postnatally rather than prenatally. Alternatively, this lack of significant difference in craniofacial form at this age could be due to developmental redundancy or compensatory growth. This is, however, consistent with the clinical phenotype of many affected individuals, in whom the onset of symptoms is recognized in early childhood rather than at birth, and underscores the importance of longitudinal assessment beginning in the perinatal period. By adulthood, cephalometric analyses of H3.3^T45I^ mice revealed skulls that appear longer and less domed compared to unaffected littermates, with a measurably enlarged synchondral growth center relative to H3.3^WT^ mice (Figure 2A-C, Supplemental Figure 2M-N). These subtle but consistent craniofacial differences parallel the variable craniofacial anomalies documented in individuals with BLBS, including the four individuals who harbor the p.T45I variant specifically (Table 1) (Bryant et al., 2020; Layo-Carris et al., 2024).

Immunohistochemical analyses of adult brain tissue revealed no significant differences between genotypes in neuronal density (NeuN), astrocyte burden (GFAP), or Purkinje cell number (Calbindin), though a sex-based difference in microglia (Iba1) was observed among H3.3^WT^ animals, with female H3.3^WT^ mice harboring significantly more microglia than their male H3.3^WT^ counterparts (Figure 2D-F). This finding may reflect known sex differences in baseline neuroimmune composition and warrants further investigation in the context of H3.3 function in the brain (Bryant et al., 2024).

A defining and clinically significant feature of BLBS is progressive neuromotor decline, and one of our primary objectives was to determine whether this trajectory is faithfully recapitulated in the H3.3^T45I^ mouse model. The longitudinal motor data presented here strongly support this assertion. During the perinatal period, H3.3^T45I^ pups demonstrated significantly delayed negative geotaxis and circle exit latency beginning at PND 12, reflecting early impairment in motor coordination and postural control (Figure 1G-H). In adulthood (12–16 weeks), rotarod performance revealed mild but statistically significant deficits in gross motor coordination in H3.3^T45I^ mice (Figure 3D). These deficits were substantially more pronounced in detailed gait analysis: male H3.3^T45I^ mice demonstrated reduced ambulatory speed, decreased cadence, increased standing time, and shortened stride length relative to H3.3^WT^ littermates (Figure 3E-J). Reduced voluntary wheel running activity during the active phase of the light-dark cycle further confirmed decreased motor output in H3.3^T45I^ mice of both sexes, though the sex-by-genotype interaction in this assay was notable (Figure 3K). The observation that fine motor phenotypes were more prominent in adult males than females was unexpected given the clinical presentation, in which no strong sex bias in motor severity has been established (Okur et al., 2021). However, by 60 weeks of rotarod deficits were present in both male and female H3.3^T45I^ mice across all nine trials, without sex-based differences (Figure 4A). Stride length remained significantly reduced in elderly H3.3^T45I^ mice (Figure 4F). Together, these data document a progressive worsening of motor performance with age in the H3.3^T45I^ model that mirrors the progressive neuromotor trajectory reported in affected individuals (Layo-Carris et al., 2024; Khazaei et al., 2023). The sex-by-age interaction observed in this model also serves as a useful reminder that phenotypic divergence between sexes may reflect differences in rate rather than trajectory of decline, a consideration that will be important in the design of future preclinical therapeutic trials.

Individuals with BLBS are frequently diagnosed with developmental delay and intellectual disability, anxiety, and/or autism spectrum disorder, and these features were also represented in the four individuals harboring the p.T45I variant (Table 1) (Layo-Carris et al., 2024). We therefore sought to determine whether the H3.3^T45I^ mouse model recapitulates relevant behavioral phenotypes beyond motor function. Notably, H3.3^T45I^ mice did not demonstrate anxiety-related behavior in the open field or elevated zero maze assays (Figure 5A-5B). Learning and memory, as assessed by spontaneous alternation in the Y-maze and novelty preference in the novel object recognition task, were intact in H3.3^T45I^ mice of both sexes (Figure 5D-5E). Similarly, all mice demonstrated appropriate social preference and social memory in the three-chamber task (Figure 5F-5G). However, a suite of more nuanced behavioral observations suggests that the H3.3^T45I^ model does capture socially and clinically meaningful phenotypes. H3.3^T45I^ mice exhibited significantly impaired nest building, failing to improve nest scores over a 24-hour period in contrast to the progressive improvement seen in H3.3^WT^ littermates (Figure 5C). Nest building is a species-typical motivated behavior and is considered an OCD-like or goal-directed behavioral measure, and deficits in this assay have been associated with prefrontal and striatal dysfunction in other neurodevelopmental mouse models (Mj, 2006; Jirkof, 2014; Kalueff et al., 2015; Shmelkov et al., 2010; Deacon et al.; Peixoto et al., 2019; Mccarthy et al., 2024). H3.3^T45I^ mice also showed significantly increased locomotion during social interaction phases of the three-chamber task (Figure 5H), and male H3.3^T45I^ mice displayed aberrant investigatory behavior characterized by increased mounting/rearing against social stimulus containers rather than the typical nose-poke investigation seen in H3.3^WT^ mice (Figure 5I). This qualitative difference in social investigation modality, combined with significantly elevated aggression in male H3.3^T45I^ mice in the resident intruder assay (Figure 5J), suggests that while the valence of social interest is intact, the expression of social behavior is dysregulated in this model. The sex specificity of the aggression phenotype is consistent with broader patterns of sex-differentiated aggression in rodent models and warrants further characterization (Aubry et al., 2022; Koolhaas et al., 2013; Takahashi, 2021). Communication phenotypes, assessed via ultrasonic vocalizations in pre-weaned pups, trended toward a reduced number of calls in H3.3^T45I^ mice but did not reach statistical significance (Supplemental Figure 4), a finding that aligns with the observation that expressive communication is variably affected in the BLBS cohort (Layo-Carris et al., 2024). Collectively, these behavior data indicate that the H3.3^T45I^ mouse model captures a subset of clinically relevant behavioral features of BLBS and that additional testing with paradigms sensitive to subtle changes in executive function and social communication may reveal further convergence.

This work builds on a growing body of functional studies that are collectively advancing the understanding of BLBS pathogenesis. Prior work demonstrated that single amino acid substitutions in H3.3 G34 modeled in mice (H3.3 p.G34R/V/W) recapitulate the mixed neurodevelopmental and neurodegenerative features of BLBS, including notable phenotypic variability (Khazaei et al., 2023). That study underscored the lesson originally demonstrated by affected individuals: there is significant phenotypic heterogeneity across the BLBS cohort, and studying multiple causal variants will be essential to identify a convergent, therapeutically targetable mechanism. More recently, the first isogenic human induced pluripotent stem cell model and three-dimensional dorsal forebrain organoid model of BLBS, harboring H3.3 p.L48R, demonstrated disruption of chromatin dynamics, neuronal differentiation, and maturation in ways that are not accessible in rodent systems (Sangree et al., 2026) . The H3.3^T45I^ mouse model presented here complements these systems. As the p.T45I variant is shared by four unrelated individuals, it is among the most commonly recurrent variants in the BLBS cohort (Bryant et al., 2020; Layo-Carris et al., 2024), making it a high-priority target for preclinical study. The robust *in vivo* behavioral and morphological phenotypes established here provide outcome measures that can serve as endpoints in future preclinical therapeutic trials. Together with prior functional models, this work reflects the importance of integrating multiple complementary model systems that collectively better represent the diversity of the BLBS community than any single model can individually (Khazaei et al., 2023; Sangree et al., 2026).

We also recognize the limitations to this work. No one model system will perfectly recapitulate every feature of a human syndrome. This goal is even less attainable for a condition in which even affected individuals harboring the same base pair substitution present with heterogeneous phenotypes (Table 1) (Bryant et al., 2020; Layo-Carris et al., 2024). For example, the sex-based dichotomy in fine motor phenotypes observed in adult H3.3^T45I^ mice was not anticipated on the basis of clinical data (Okur et al., 2021), and the molecular basis of this difference remains to be determined. Similarly, structural brain abnormalities are variably present in affected individuals (Table 1) (Okur et al., 2021; Layo-Carris et al., 2024), yet gross neuronal and glial cell type composition appeared largely intact at the adult time points examined here, suggesting that the relevant neuropathological differences may lie in connectivity, synaptic function, epigenetic state, or transcript regulation rather than in cell numbers alone (Bryant et al., 2024; Xia and Jiao, 2017). Future work integrating transcriptomic and chromatin accessibility analyses will be essential to characterize the molecular underpinnings of the behavioral and morphological phenotypes described here. Nonetheless, we feel that identifying and embracing this heterogeneity at this stage is a significant asset as we move forward toward therapy development. A robust body of literature demonstrates that failure to account for genetic and phenotypic variability contributes to the staggering statistic that ∼90% of clinical trials fail (Woodward et al., 2022; Latapiat et al., 2023; Minikel et al., 2024; Razuvayevskaya et al., 2024). Thus, integrating the strengths of multiple model systems available for BLBS translational drug discovery efforts, that when combined better reflect the diverse community than any one model individually, enhances confidence in the robustness and replicability of future preclinical trials (Khazaei et al., 2023; Sangree et al., 2026).

In conclusion, we present a comprehensive characterization of a novel mouse model of BLBS harboring the recurrent H3.3 p.T45I missense variant. The developmental delays, progressive motor decline, and selected behavioral changes are reminiscent of symptoms observed in individuals with BLBS, including those harboring this specific variant. Thus, providing a useful tool for the development of future therapies. The non-invasive, clinically translatable outcome measures established here, including growth trajectories, gait parameters, balance/coordination performance, and behavioral endpoints, are well-suited, quantifiable outputs to be investigated in future preclinical intervention studies. As disease-modifying therapies begin to fundamentally alter the clinical trajectory for children with other pediatric neurodegenerative conditions (Mendell et al., 2017; Mercuri et al., 2018; Mendell et al., 2025; Gangji et al., 2025; Scott et al., 2023; Musunuru et al., 2025), this work contributes an essential preclinical resource toward achieving the same goal for the BLBS community.

## Acknowledgements

The authors would like to express appreciation for the University of Pennsylvania Perelman School of Medicine CRISPR-Cas9 Mouse Targeting Core Facility (RRID: SCR_022378), with special acknowledgement of Dr. Leonel D. Joannas. We also thank the Neurobehavior Testing Core at UPenn/ITMAT P50HD105354 for assistance with performing the behavior procedures. We would also like to acknowledge the tremendous work of the rare disease community in highlighting the need for and utility of these preclinical models. This work has shaped our understanding of translational research, and we hope that this work will someday be transformational for the BLBS community and others in the rare disease space.

## Author Contributions

*Conceptualization*: DELC, ELD, EEL, LMB, EJKB

*Methodology:* DELC, ELD, EEL, AKS, BC, EMG, KJA, DN, WTO, LMB

*Software*: DELC, ELD, EEL

*Validation*: DELC, ELD, EEL, AKS, BC, MH, SMS, KTW, HME, EMG, XMW, EW, KJA, DN

*Formal Analysis*: DELC, ELD, EEL, MH, HME

*Investigation:* DELC, ELD, EEL, AKS, BC, MH, SMS, KTW, HME, EMG, XMW, EW, KJA, DN, LMB

*Resources*: EJKB, WTO

*Data Curation*: DELC, ELD, EEL, BC, WTO

*Writing – Original Draft Preparation*: DELC, ELD, EEL

*Writing – Review and Editing:* DELC, ELD, EEL, AKS, BC, MH, SMS, KTW, HME, EMG, XMW, EEW, KJA, DN, WTO, LMB, EJKB

*Visualization:* DELC, ELD, EEL *Supervision*: DELC, EJKB, WTO *Project Administration:* EJKB

*Funding Acquisition*: DELC, ELD, EEL, LMB, EJKB

## Materials and Methods

### Model

The *H3-3A* p.T45I mouse model was created by the CRISPR-Cas9 Mouse Targeting Core at the University of Pennsylvania on the C57BL/6J background. The *H3-3A* sequence is homologous between mice and human sequences, so there was no effort to humanize the mouse. CRISPR-Cas9 technology was used to replace c.137C>T to produce the p.T45I variant. The Children’s Hospital of Philadelphia Institutional Animal Care and Use Committee (IACUC) approved this study. Both male and female mice were included in all experiments to account for sex as a biological variable.

### Skull Form Analysis

H3.3^T45I^ mice and age-matched wildtype littermate controls were collected at P0 and adult time points. After euthanasia with CO_2_, cervical dislocation, and decapitation, heads were stored in paraformaldehyde for 48 hours, then transitioned to 70% ethanol at 4°C until µCT images were obtained. Skulls were scanned with a Scanco µCT35 (SCANCO Medical AG, Switzerland, Penn Center for Musculoskeletal Disorders MicroCT Imaging Core, Perelman School of Medicine, University of Pennsylvania) at a resolution of 15µm. Skulls were reconstructed with Scanco software and analysis of images was performed with 3DSlicer and MeshInspector (V5.2.2, Slicer.org) (Fedorov et al., 2012).

Threshold settings were optimized to visualize only bone volume. Virtual endocasts of the skull were created to compare skull volumes by using 3DSlicer Software and the Wrap Solidify extension. Cephalometric quantifications from these images were measured from the 3D skull models including length, width, height, and endocranial volume. These measurements were inferred using previously published landmarks (Durham et al., 2019).

### Histology

Mice were perfused with 1X PBS solution followed by 4% paraformaldehyde. Brains were then carefully dissected from H3.3^T45I^ mice and age matched wildtype littermate controls. Brains were flash frozen in liquid nitrogen, embedded in OCT compound, allowed to equilibrate in the cryostat (Microm HM 505E, CHOP Pathology Core) for at least 30 minutes before sectioning. Sections were cut at 16um using a high-profile blade (Leica Biosystems 14035838926) and mounted on Colorfrost Plus Microscope Slides (Fisher Scientific 12-550-17). Slides were stored at −80C until staining when they were allowed to come to room temperature. Hydrophobic barriers were drawn around sections, and proper blocking and incubation with primary (Table 2) and appropriate secondary antibodies (dilution 1:1000) followed.

**Table 2:** Primary Antibodies for Immunohistochemical Analysis of Murine Brain components.

| Antibody | Manufacturer | Dilution |
| --- | --- | --- |
| GFAP | NOVUS (NB120-16997) | 1:250 |
| Calb1 | Antibodies Inc. (75-448) | 1:250 |
| Iba1 | Antibodies Inc. (IBA1-0100) | 1:250 |
| Oligo2 | Antibodies Inc. (OLIG2-0100) | 1:500 |
| GFAP | Antibodies Inc. (N206A/8) | 1:500 |
| NeuN | Millipore Sigma (MAB377X) | 1:50 |

### Behaviors

Developmental milestones were collected every other day between post-natal day 2 and 20. At least 10 litters were tested. All researchers were blinded to genotype. Weight, length, fur development, pinnae detachment, eyelid opening, and incisor eruption were assessed based on standard protocol (Musi et al., 1994). Pups were assessed for surface righting responses, negative geotaxis, T-bar suspension, circle motor movement, and horizontal/vertical screen suspension (Lynch et al., 2020).

Behavior data were collected on adult mice (12-16 weeks old) and aged mice (60 weeks). Motor, cognitive and social behavior assays were performed across multiple cohorts of mice. Adult mice were assessed for motor output activity with an open field arena, three-day accelerating rotarod, the CatWalkXT gait analysis system, and wheel running. Anxiety-related and social behavior were assessed through an elevated zero maze, center activity in an open field, nest building, resident intruder assay, and the three-chamber social preference. Learning and memory procedures include spontaneous alternations in a Y-maze, , novel object recognition, and a social memory phase performed in the three-chamber arena (Lynch et al., 2020).

### Statistics

Comparisons were made between H3.3^WT^ and H3.3^T45I^ mice at each time point using GraphPad PRISM 10 software and Two sample T-tests. Sub analyses were conducted by sex. For more complex assessments over time and across multiple groups (male and female for each genotype), Kruskal Wallis non-parametric assessments were used, as data lacked normal distribution. Data are presented as mean ± standard error mean. p≤0.05 was considered significant. No specific power analysis was conducted for these assessments, however an n of 8 was determined to be appropriate for other neurodegenerative disease mouse models, therefore our assessments include at least 8 individuals per genotype (Lynch et al., 2020).

---

**Supplemental Figure 1.**
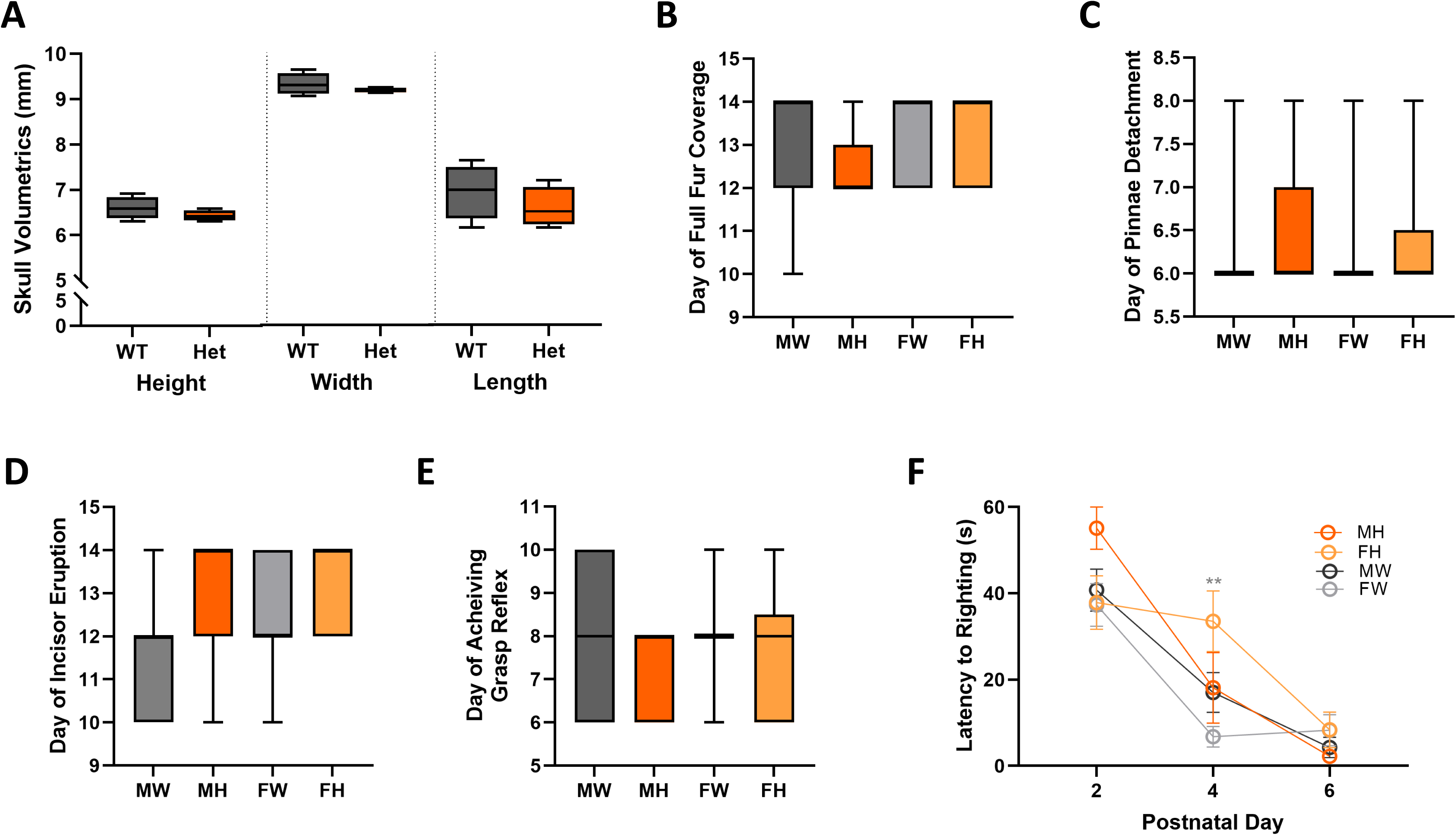
H3.3 pups show few differences from H3.3 littermates in perinatal skull form or non-motor developmental milestones. A–F. Male H3.3^WT^ data is black/dark gray, female H3.3^WT^ data is light grey, male H3.3^T45I^ data is orange, and female H3.3^T45I^ data is pale orange. A. Cephalometric measure of three-dimensional µCT reconstructions of H3.3^WT^ and H3.3^T45I^ skulls at P0 show no differences in skull height, width, or length indicating potentially subtle brain differences if any between genotypes. n=4 B. No statistically significant differences were observed between H3.3^T45I^ pups and H3.3^WT^ littermates in day of fur eruption. C. No statistically significant differences were observed between H3.3^T45I^ pups and H3.3^WT^ littermates in day of pinnae detachment. D. No statistically significant differences were observed between H3.3^T45I^ pups and H3.3^WT^ littermates in day of incisor eruption. E. No statistically significant differences were observed between H3.3^T45I^ pups and H3.3^WT^ littermates in day of achieving grasp reflex. F. No statistically significant differences were observed between H3.3^T45I^ pups and H3.3^WT^ littermates in latency to right in the surface righting assay. n ≥ 8 for assessments other than Micro-CT. MW = male WT; MH = male Het; FW = female WT; FH = female Het.

**Supplemental Figure 2.**
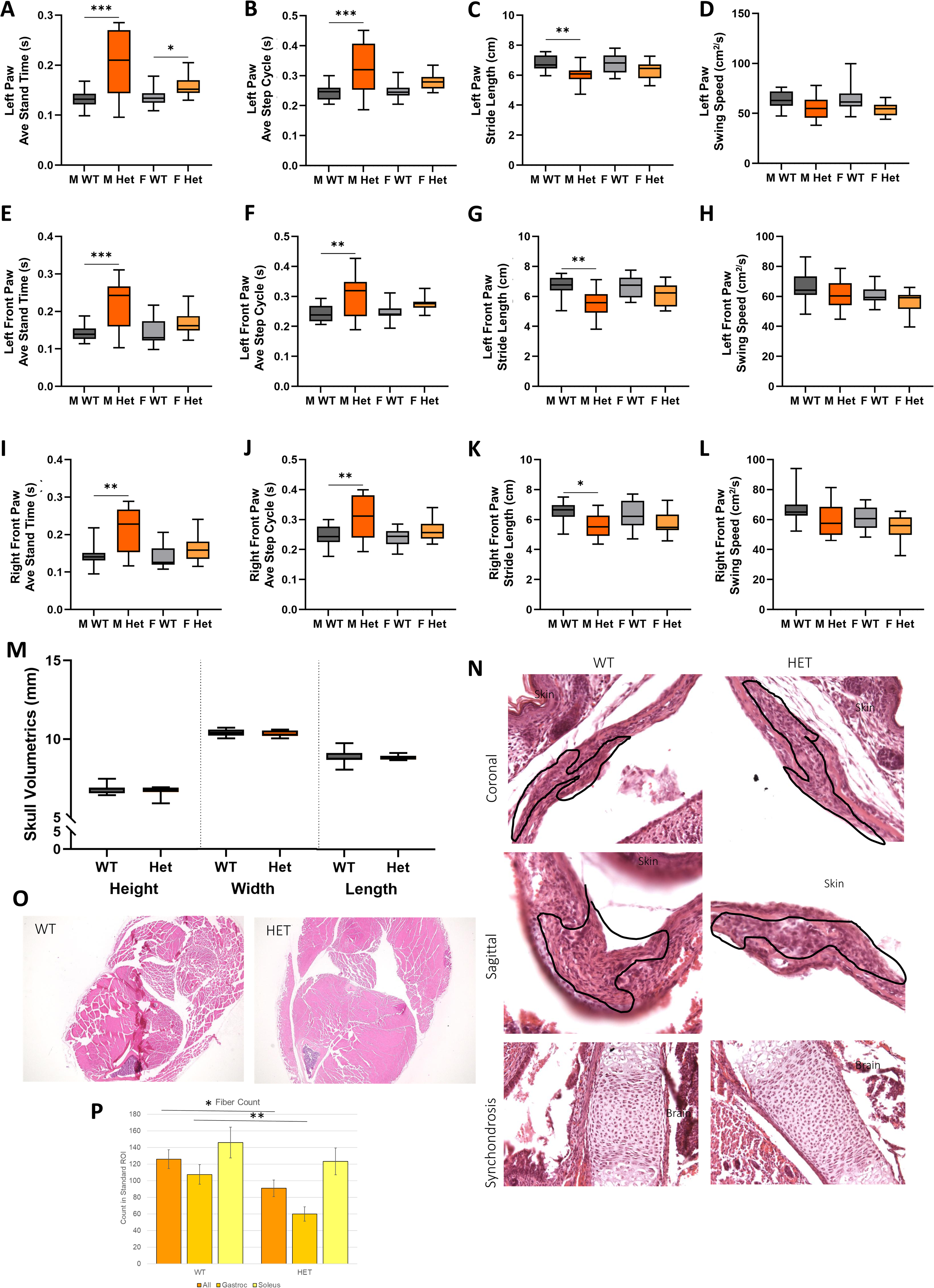
Comprehensive CatWalkXT gait metrics, adult craniofacial morphometrics, and lower limb histology in adult H3.3^T45I^ mice. A–L. Male H3.3^WT^ data is black/dark gray, female H3.3^WT^ data is light grey, male H3.3^T45I^ data is orange, and female H3.3^T45I^ data is pale orange. A–L. Complete Noldus CatWalk gait analysis data for all four paws in adult (12–16 week) H3.3^T45I^ mice and H3.3^WT^ littermates. Parameters shown include average speed, cadence, stand time, step cycle, stride length, and swing speed for left front, right front, left hind, and right hind paws. Male H3.3^T45I^ mice show consistent deficits in ambulatory speed, cadence, stride length, and step cycle parameters across paws, while female H3.3^T45I^ mice do not show statistically significant differences from H3.3^WT^ littermates. Right paw data for representative parameters are shown in Figure 3. M. Cephalometric measure of three-dimensional µCT reconstructions of H3.3^WT^ and H3.3^T45I^ skulls show no differences in skull height, width, or length indicating potentially subtle brain differences if any between genotypes. H3.3^T45I^ skulls appear longer and less domed compared to H3.3^WT^ littermates. N. Representative hematoxylin and eosin-stained histological images highlighting the coronal suture (top) sagittal suture (middle) and sphenooccipital synchondrosis (SOS, bottom) in adult H3.3^WT^ (left) and H3.3^T45I^ (right) mice. H3.3^T45I^ mice show a measurably larger SOS growth center compared to H3.3^WT^ littermates. O–P. Representative hematoxylin and eosin-stained histological sections of lower limb skeletal muscle from adult H3.3^T45I^ (right) and H3.3^WT^ (left) littermates. Subtle differences in muscle fiber morphology are observed in H3.3T45I mice compared to unaffected controls that are statistically significant when fibers are counted (p=0.017), particularly in the gastrocnemius muscle (p=0.005). n≥8 MW = male WT; MH = male Het; FW = female WT; FH = female Het. *p≤0.05, ** p≤0.01 *** p≤0.001.

**Supplemental Figure 3.**
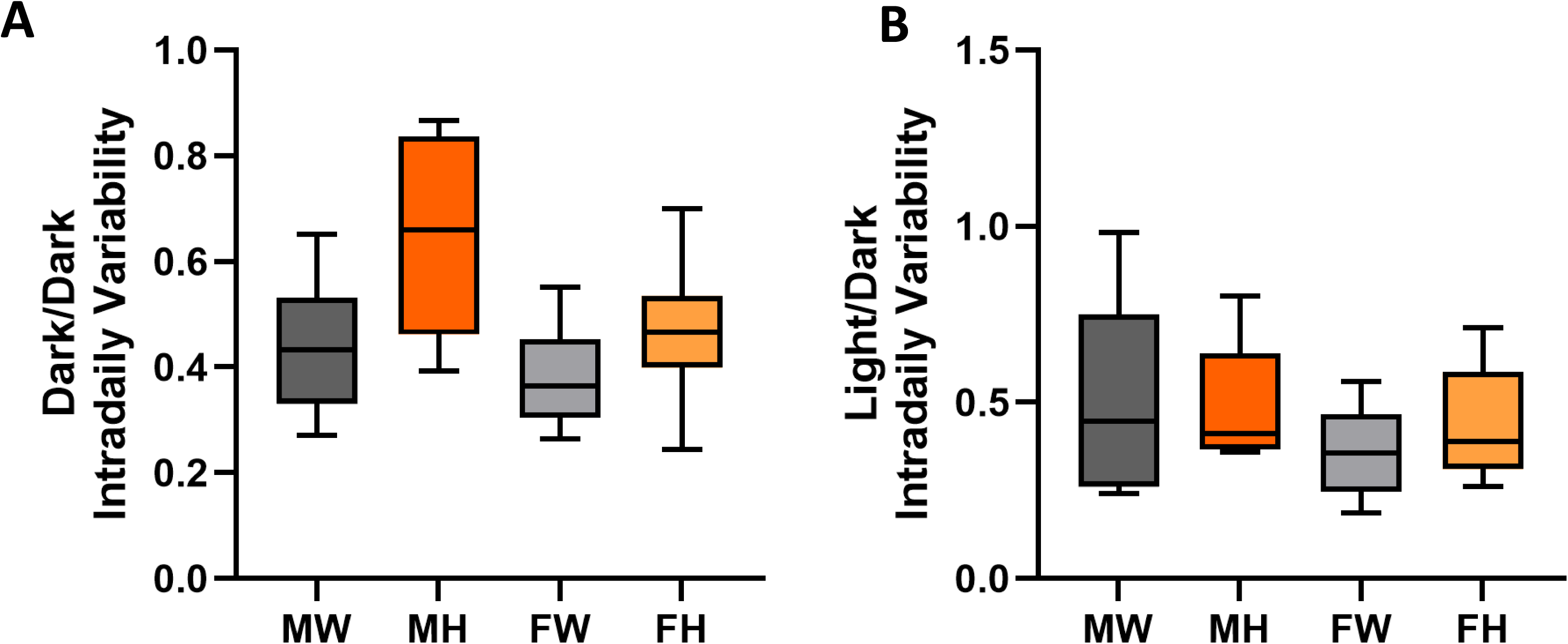
Complete light/ dark wheel running cycle assessment for adult H3.3 mice. A–B. Wheel running was assessed over a 24-hour period to investigate genotype- and sex-driven differences at the intersection of motor development and circadian rhythm regulation. ZT0–ZT12 reflects the light phase and ZT12–ZT24 the dark phase. A. Dark/Dark Intradaily Variability of wheel running across the dark phase. B. Light/Dark Intradaily Variability of wheel running across the light phase. No statistically significant shifts in Zeitgeber time were observed between H3.3^T45I^ mice and H3.3^WT^ littermates. Male H3.3^WT^ data is black/dark gray, female H3.3^WT^ data is light grey, male H3.3^T45I^ data is orange, and female H3.3^T45I^ data is pale orange. MW = male WT; MH = male Het; FW = female WT; FH = female Het. n≥8

**Supplemental Figure 4.**
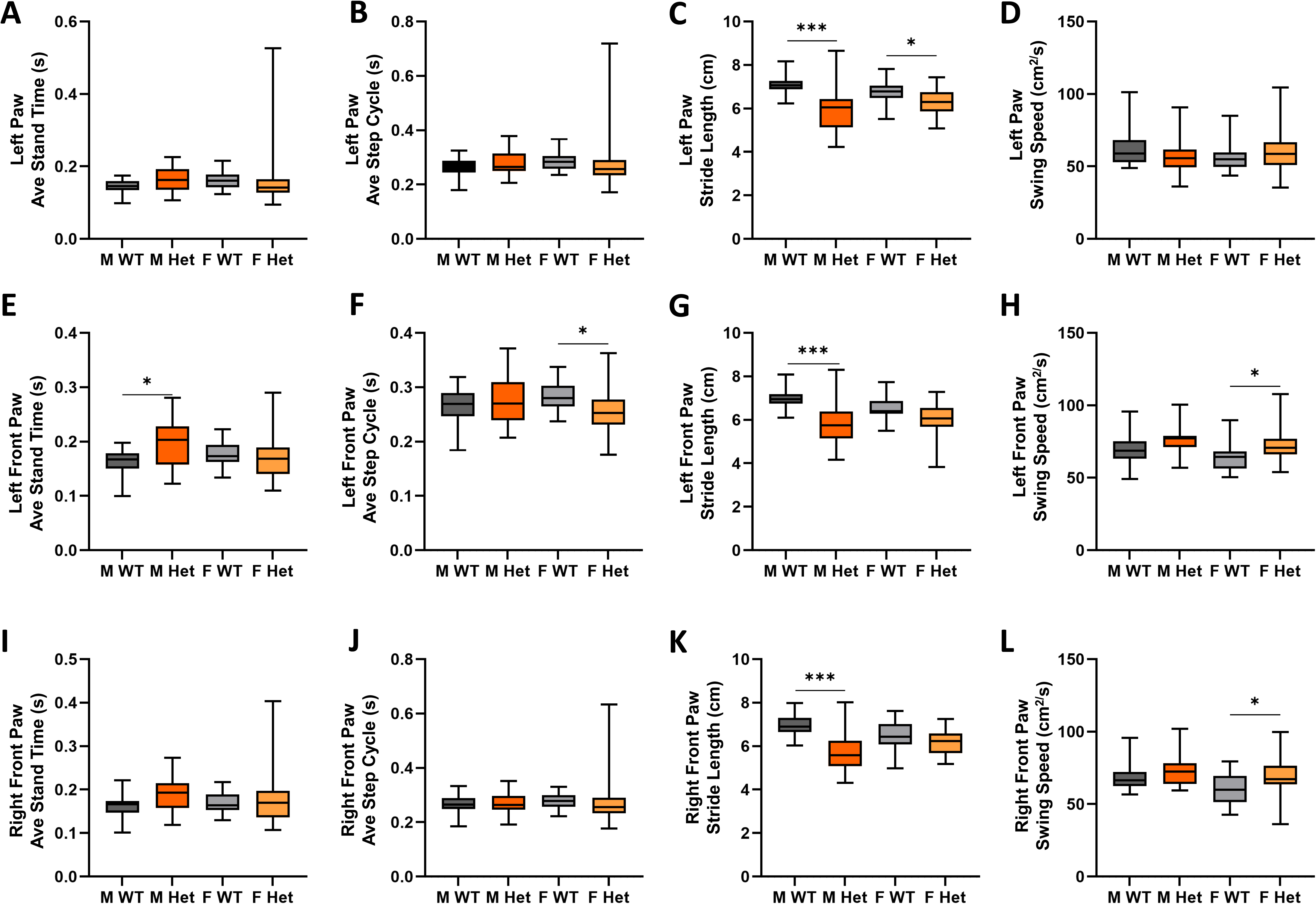
Complete CatWalkXT gait metrics for all paws in elderly H3.3 mice. A–L. Male H3.3^WT^ data is black/dark gray, female H3.3^WT^ data is light grey, male H3.3^T45I^ data is orange, and female H3.3^T45I^ data is pale orange. A–L. Complete Noldus CatWalk gait analysis data for all four paws in elderly (≥60 week) H3.3^T45I^ mice and H3.3^WT^ littermates. Parameters shown include average speed, cadence, stand time, step cycle, stride length, and swing speed for left front, right front, left hind, and right hind paws. Compared to the adult cohort, elderly H3.3^T45I^ mice show greater within-group variation and fewer statistically significant individual gait parameters, with the exception of stride length, which remains significantly reduced in both male (p≤0.001) and female (p=0.028) H3.3^T45I^ mice relative to H3.3^WT^ littermates. Unlike in the adult cohort, no sex-driven differences in gait metrics are observed at this timepoint. Right paw data for representative parameters are shown in Figure 4. n≥8. MW = male WT; MH = male Het; FW = female WT; FH = female Het. *p≤0.05, ** p≤0.01 *** p≤0.001.

**Supplemental Figure 5.**
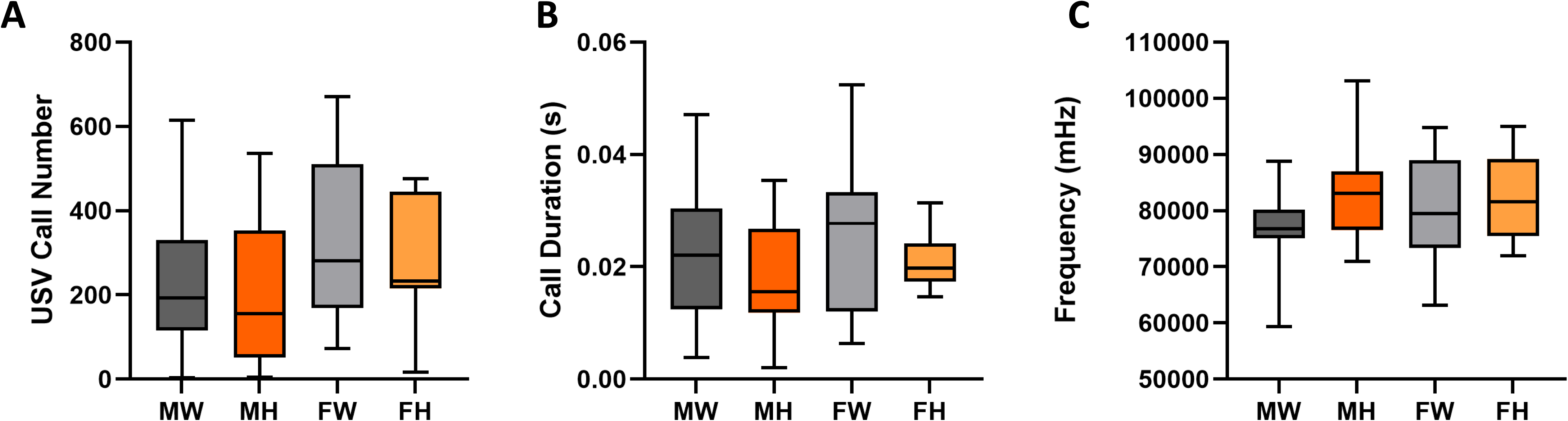
Ultrasonic vocalizations in pre-weaned H3.3T45I pups trend toward reduction but do not reach statistical significance. A–C. Male H3.3^WT^ data is black/dark gray, female H3.3^WT^ data is light grey, male H3.3^T45I^ data is orange, and female H3.3^T45I^ data is pale orange. A. Total number of ultrasonic vocalization (USV) calls recorded from pre-weaned H3.3^T45I^ pups and H3.3^WT^ littermates. Male and female H3.3^T45I^ pups trended toward fewer USV calls compared to unaffected littermates, but this difference did not reach statistical significance. B. Total duration of USV calls recorded from pre-weaned H3.3^T45I^ pups and H3.3^WT^ littermates. Male and female H3.3^T45I^ pups trended toward shorter total USV call duration compared to unaffected littermates, but this difference did not reach statistical significance. C. USV call frequency (kHz) recorded from pre-weaned H3.3^T45I^ pups and H3.3^WT^ littermates. No clear sex- or genotype-driven differences in USV frequency were observed. n≥8. MW = male WT; MH = male Het; FW = female WT; FH = female Het.

